# PatientPheX: A Full-Text Biomedical Corpus for Patient-Specific Phenotype Information Extraction

**DOI:** 10.64898/2026.10.07.757319

**Authors:** Xuanlyu Zhu, Ling Luo, Zhijun Wang, Chentao Huang, Yunxiang Li, Yang Zhong, Tengxiao Lv, Juntao Li, Chao Liu, Feng Tian, Yuanyuan Sun, Jian Wang, Hongfei Lin

**Affiliations:** School of Computer Science and Technology, Dalian University of Technology, Dalian 116024, China

**Keywords:** patient-specific phenotype extraction, annotated biomedical corpus, named entity recognition, relation extraction, biomedical full text

## Abstract

Patient-specific phenotype information extraction identifiers and normalizes phenotype entities and links them with individual patients, supporting rare disease research and genotype-phenotype analysis. However, existing datasets rarely combine full-text biomedical content with detailed annotations of patients, phenotypes, related entities, and their associations. In this work, we present PatientPheX, a corpus for patient-specific phenotype information extraction comprising 200 full-text articles, primarily case reports and case series. The corpus provides mention annotations and normalization for seven biomedical entity types, together with explicit associations between patients and entities across sentences, paragraphs, and sections. Negated findings and composite phenotype mentions are also annotated. Corpus construction combines automated pre-annotation, large language model (LLM)-assisted verification, and manual review and adjudication. Benchmarks cover entity recognition and normalization and patient–phenotype association extraction using gold-standard and predicted entities. The overall document-level entity F1 was 86.95%, while phenotype F1 was 79.33%. The best relation extraction macro-F1 decreased from 84.14% with gold-standard entities to 54.98% with predicted entities, highlighting the combined challenges of phenotype extraction and patient attribution. PatientPheX provides a resource for developing and evaluating patient-centered information extraction methods and supports the construction of structured patient-level knowledge resources for phenotype-driven biomedical research.

## 1 Introduction

Phenotypes are observable or measurable characteristics of an individual. In clinical contexts, phenotypic abnormalities include signs, symptoms, and abnormal findings from physical examinations, laboratory tests, and medical imaging. Detailed and standardized phenotype information plays an important role in rare disease diagnosis, disease characterization, and the investigation of genotype-phenotype relations [1,2]. The Human Phenotype Ontology (HPO) provides a standardized vocabulary for describing phenotypic abnormalities, enabling phenotype information to be compared and analyzed across patients and data sources [2]. Biomedical literature, particularly case reports and case series, contains rich descriptions of individual patients and their clinical findings. However, these descriptions are largely recorded in unstructured text, which limits their systematic retrieval and integration. Patient-specific phenotype information extraction (IE) addresses this challenge by identifying phenotype mentions, normalizing them to standardized concepts, and linking them to individual patients.

Patient-specific phenotype IE involves three sub-tasks: named entity recognition (NER), which identifies phenotype and related biomedical mentions and their textual spans; named entity normalization (NEN), which maps these mentions to standardized concepts (e.g., HPO terms); and relation extraction (RE), which associates the extracted entities with specific patients. Related biomedical entities, including diseases, genes, variants, and chemicals, provide diagnostic, genetic, and therapeutic context for interpreting a patient’s phenotype profile. Accurate patient attribution is essential because a single article may describe multiple patients, relatives, or controls. Furthermore, negated findings and general background information should also be distinguished from findings actually observed in a patient [3]. These distinctions are particularly challenging in full-text articles, where patient references and relevant findings may be distributed across sentences, paragraphs, and sections.

Several annotated corpora have been developed to support phenotype recognition, normalization, and patient-level representation [4-9]. Nevertheless, existing resources provide limited support for extracting rich, patient-specific information from full-text biomedical literature. Corpora with mention-level phenotype annotations commonly focus on abstracts, short case descriptions, or clinical observations, providing limited context for identifying and associating findings distributed across an entire article. Other resources provide document-level or patient-level HPO profiles but do not consistently preserve the underlying text spans or explicit associations between phenotype mentions and individual patients. Few resources therefore combine full-text content, mention-level annotations, standardized biomedical concepts, and associations linking patients to phenotypes and related biomedical entities. This lack of integrated annotations limits the development and evaluation of methods that transform information dispersed throughout an article into structured patient-specific representations.

To address this limitation, we present PatientPheX, a corpus of 200 full-text biomedical articles, primarily case reports and case series, for patient-specific phenotype IE. Corpus construction combines automated pre-annotation, large language model (LLM)-assisted verification, and manual review and adjudication. PatientPheX supports the recognition and normalization of phenotypes and related biomedical entities, together with their attribution to individual patients. Its main features are summarized as follows:

i. Integrated phenotype and biomedical entity annotations. PatientPheX includes mention-level annotations for seven entity types: phenotype, disease, gene, chemical, variant, cell line, and species. These mentions are normalized to the ontology concepts, while related entities provide diagnostic, genetic, therapeutic, and biological context for interpreting individual patients’ clinical findings.
ii. Annotations of complex clinical phenotype expressions. The corpus includes negated and composite phenotype mentions, supporting the distinction between present and absent findings and the representation of expressions that encode multiple phenotype concepts.
iii. Document-level extraction of patient-specific information. Explicit associations link phenotypes and related biomedical entities to individual patients across sentences, paragraphs, and sections. These annotations support the development and evaluation of methods that resolve patient attribution throughout an article, including cases involving multiple patients or family members, and organize dispersed findings into structured patient-specific profiles.

To assess the utility of PatientPheX and characterize its extraction challenges, we establish baselines for entity recognition and normalization across the seven entity types and for patient-phenotype association extraction. These experiments evaluate the performance of existing methods in identifying biomedical concepts and assigning phenotypes to the correct patients. We also organized a shared evaluation task (https://github.com/DUTIR-BioNLP/PatientPheX), focusing on phenotype recognition, HPO normalization, and patient-phenotype association extraction. The baseline experiments and shared evaluation provide reference results for developing methods that extract and organize patient-specific phenotype information from full-text biomedical literature.

## 2 Related Work

Several corpora have been developed to support phenotype concept recognition and patient-level phenotype representation. Table 1 compares representative annotation corpora and structured knowledge resources in terms of source material, size, annotation scope, and patient-related information. GSC and its updated version, GSC 2024, provide mention-level phenotype annotations linked to HPO concepts in biomedical abstracts [4,5]. ID-68 extends phenotype annotation to clinical case descriptions [6], while BioCreative VIII Task 3 (BC8-T3) focuses on observations from dysmorphology physical examination observations and includes negated, overlapping, discontinuous, and descriptive phenotype mentions [7]. These corpora support the development and evaluation of phenotype concept recognition methods. However, their focus on abstracts, short case descriptions, or individual clinical observations limits their use for evaluating extraction and patient attribution across full-text articles.

**Table 1.** Comparison of phenotype annotation datasets.

| Dataset | Text type | Size | Span | HPO normalization | Negation | Composition | Relation annotation |
| --- | --- | --- | --- | --- | --- | --- | --- |
| GSC 2024 [5] | PubMed abstracts | 228 abstracts | ✓ | ✓ | – | – | – |
| ID-68 [6] | Clinical case descriptions | 68 cases | ✓ | ✓ | – | – | – |
| BC8-T3 [7] | Dysmorphology observations | 3,136 observations | ✓ | ✓ | ✓ | ✓ | – |
| JAX [8] | PMC full-text articles | 131 articles | – | ✓ | – | – | – |
| CSC [9] | Published case reports | 112 cases | – | ✓ | – | – | – |
| Liu et al. [3] | Clinical notes | 50 notes | ✓ | ✓ | ✓ | – | Patient-specific status |
| Phenopacket Store [10] | Structured cases curated from literature | 6,668 individuals | – | ✓ | ✓ | – | Patient-level structured associations |
| FPKG [11] | Structured knowledge curated from biomedical literature | 509 source articles; 6,143 nodes; 19,282 relations | – | Node-level HPO IDs | – | – | Curated graph relations, including patient-related associations |
| PatientPheX (Ours) | Full-text case reports and case series | 200 articles | ✓ | ✓ | ✓ | ✓ | Patient–entity association |

Note: ✓ indicates that the annotation feature is included; – indicates that it is not reported as a dedicated annotation feature in this comparison.

Other resources support phenotype representation at the document or patient level. The JAX corpus provides document-level HPO annotations for full-text biomedical articles [8], while the Case Study Cohort (CSC) associates HPO concepts with published clinical cases to evaluate automated deep phenotyping methods [9]. Liu et al. annotated patient-specific phenotypes in clinical notes, distinguishing findings associated with the target patient from negated findings and family-history information [3]. Although these resources support phenotype representation beyond individual mentions, they address different aspects of patient attribution. JAX and CSC primarily provide document-level or case-level HPO profiles rather than explicit links between textual mentions and individual patients. The clinical-note corpus of Liu et al. focuses on a predefined target patient rather than attribution among multiple individuals described within the same document.

Structured knowledge resources further organize phenotypes and related clinical information around individual patients. The Phenopacket Store represents literature-derived phenotypes, diagnoses, genetic variants, and other clinical information using standardized Global Alliance for Genomics and Health (GA4GH) Phenopackets [10]. Song et al. constructed a Facial Phenotype Knowledge Graph (FPKG) from biomedical publications, integrating facial phenotypes, genes, diseases, variants, patients, and source articles to support knowledge retrieval and rare genetic disease diagnosis [11]. These resources capture patient-level biomedical associations, but their primary purpose is structured knowledge representation rather than text annotation for evaluating mention recognition and patient attribution.

PatientPheX complements these resources by integrating full-text mention annotations, biomedical entity normalization, annotations of negated and composite phenotype mentions, and explicit associations between patients and biomedical entities. This combination supports the evaluation of both phenotype extraction and document-level patient attribution, enabling findings distributed throughout an article to be organized into structured patient-specific profiles.

## 3 Methods

The construction of PatientPheX was guided by an annotation schema and followed four stages: data collection, automated pre-annotation and LLM-assisted verification, manual annotation and adjudication, and post-processing and quality control (Figure 1). This workflow combined automated assistance with independent human annotation to reduce annotation workload while maintaining consistent entity annotations and patient attribution.

**Figure 1.**
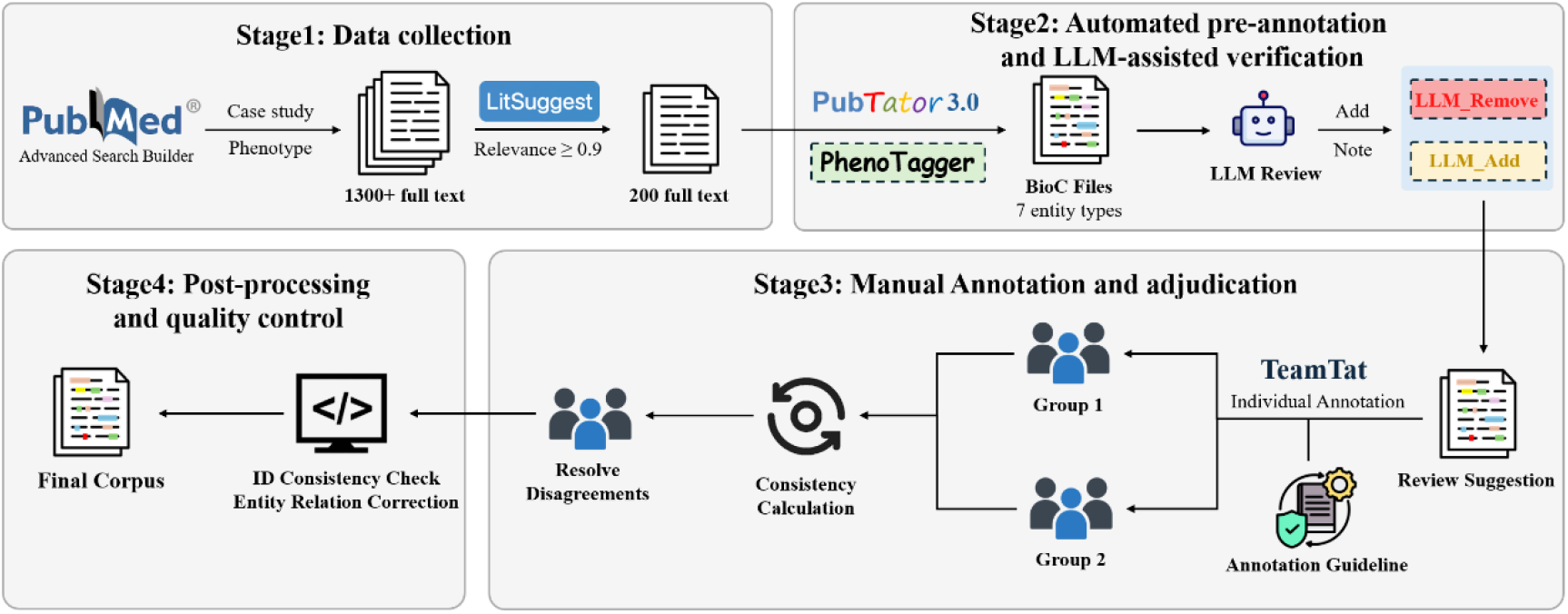
Overview of the PatientPheX corpus construction workflow.

### 3.1 Annotation scope and schema

Annotation covered the main narrative text of each article, including figure and table captions that formed part of the narrative. The internal contents of figures and tables, reference lists, acknowledgments, abbreviation lists, and other non-narrative sections were excluded. This scope maintained a consistent textual basis for annotation and excluded structurally complex content that could not be reliably represented using linear character offsets.

The annotation schema included seven entity types: (1) Phenotype: observable or measurable abnormalities, including signs, symptoms, and abnormal clinical findings; (2) Disease: named diseases, disorders, syndromes, and diagnoses; (3) Gene: genes and gene products, including proteins and messenger RNAs; (4) Chemical: drugs and other chemical substances; (5) Variant: genomic and protein sequence alterations, including substitutions, deletions, insertions, and other variant types; (6) CellLine: established cell lines used in biomedical research; (7) Species: mentions of human-including patients, probands, family members, and controls-as well as animals, microorganisms, and other organisms. Phenotype was the primary entity type, with additional annotations for negated findings and composite phenotype mentions. The remaining six categories were adapted from BioRED [13] and PubTator [14] to promote compatibility with existing biomedical corpora and normalization tools. The Species category additionally encompassed human references relevant to patient identification.

For relation annotation, entities were linked to individual patients using a single relation type of *association*, when supported by textual evidence as shown in Figure 2. These associations could extend across sentences, paragraphs, and sections, enabling findings distributed throughout an article to be organized by patient. Detailed entity definitions, normalization rules, relation criteria, and conventions for annotating complex expressions are provided in the annotation guidelines.

**Figure 2.**
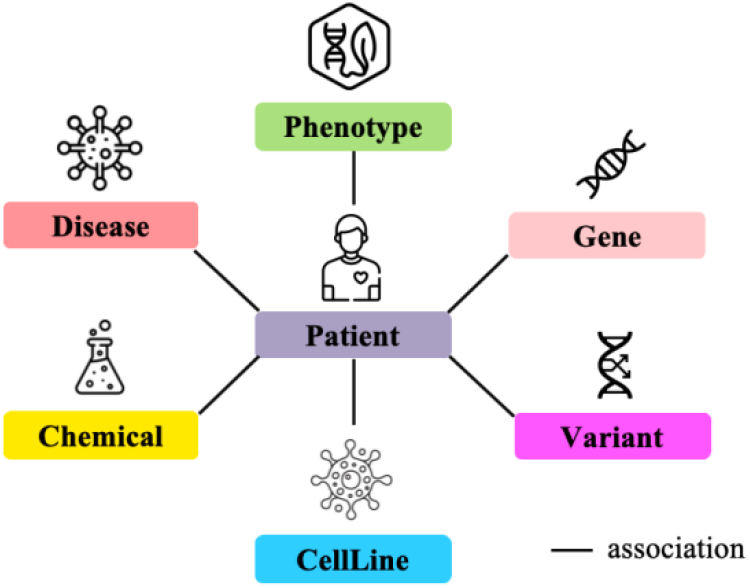
Entity types and patient-entity associations represented in PatientPheX.

### 3.2 Data collection

We focused on full-text case reports and case series indexed in PMC because they describe individual patients and provide detailed information on clinical manifestations, diagnoses, genetic findings, treatments, and follow-up outcomes. These articles offer suitable source material for studying the attribution of biomedical findings to specific patients. We used articles from the JAX corpus [8] as the initial collection and expanded it using LitSuggest [12] to identify similar, relevant publications. Additional articles were retained if they had a relevance score above 0.9, accessible full text, and a valid PubMed Central identifier (PMCID). The resulting collection provided the source material for selecting the 200 articles included in PatientPheX.

### 3.3 Automated pre-annotation and LLM-assisted verification

Using the selected PMCIDs, we retrieved full-text articles in BioC XML format through the PubTator API, together with pre-annotations for Disease, Gene, Chemical, Variant, CellLine, and Species. Phenotype mentions were identified and normalized using PhenoTagger [8], trained using the HPO released version v2026-06-23.

Before manual annotation, an LLM reviewed the source text and automated annotations to flag potential errors in mention boundaries, entity types, and normalized identifiers. It also identified potentially missing entities and ambiguous patient attribution. Suggested additions, deletions, and corrections were submitted for human review rather than incorporated directly into the gold-standard corpus.

### 3.4 Manual annotation and adjudication

Manual annotation was performed by eleven researchers with experience in biomedical natural language processing, including one professor and ten post-graduate students. The annotators were organized into five pairs and used TeamTat [15] for collaborative annotation. Prior to formal annotation, a preliminary guideline covering entity definitions, mention boundary rules, normalization, patient identification, negation, composite entities, and relation criteria was developed. All annotators participated in a pilot annotation exercise involving 20 full-text articles. Ambiguities concerning phenotype boundaries, distinctions between diseases and phenotypes, overlapping and composite mentions, patient references, and attribution among multiple patients were discussed collectively. The guidelines were refined to establish consistent annotation rules.

During formal annotation, each article was independently annotated by two annotators using the pre-annotations as a starting point. Annotators corrected entity spans, types, and normalized identifiers, removed erroneous annotations, and added missing entity mentions. They also identified patient mentions, resolved references to the same individual, and annotated associations between specific patients and biomedical entities based on evidence throughout the article. Each pair then discussed discrepancies, with unresolved or particularly ambiguous cases referred to the senior annotator for adjudication. The resulting consensus annotations formed the gold-standard corpus.

### 3.5 Post-processing and quality control

Following adjudication, programmatic checks identified formatting and consistency errors, including invalid character offsets, mismatches between annotated spans and mention lengths, duplicate or inconsistent entity identifiers, nonstandard patient identifiers, invalid relation references, and annotations in excluded sections. An LLM-based agent additionally examined textual evidence and contextual consistency to flag potentially incorrect entity annotations and patient–entity associations. The agent did not modify the annotations directly. Human annotators reviewed all flagged cases, and only manually confirmed corrections were incorporated into the final corpus.

### 3.6 Benchmark methods

We benchmarked existing biomedical information extraction methods on PatientPheX to assess their performance and identify challenges in entity recognition, normalization, and patient–phenotype association extraction. For entity recognition and normalization, PubTator [14] was used to identify six biomedical entity types, including Disease, Gene, Chemical, Variant, CellLine, and Species, while PhenoTagger [8] was used to recognize phenotype mentions and normalize them to HPO concepts. Both tools were applied directly without additional training on PatientPheX.

For patient-phenotype RE, we constructed a pipeline that linked phenotype candidates identified and normalized by PhenoTagger to individual patients. We evaluated two supervised document-level RE models, ATLOP [16] and GLiM [17], and three LLMs: Qwen3-8B, GLM-5.1, and GPT-5.4-mini. ATLOP and GLiM were trained on the PatientPheX training set. ATLOP represented patients and phenotypes as document-level entities and classified their relations using entity-aware contextual representations. GLiM additionally modeled interactions among patients, phenotypes, and entity pairs using a graph Transformer.

The three LLMs used a shared two-stage prompting strategy selected on the development set, without fine-tuning on PatientPheX. In the first stage, each article was divided into text segments. The model identified mentions of predefined patients, resolved patient references within each segment, and proposed associations with phenotype candidates in the local context. In the second stage, the model received the full section context and preliminary associations to refine patient attribution, resolve references across sentences and segments, and identify potentially missed associations. Predictions were then aggregated across the article and deduplicated to produce a set of phenotype associations for each patient. All three LLMs were evaluated in a zero-shot setting using the same phenotype candidates and prompting strategy to facilitate comparison of their patient attribution performance.

We evaluated entity recognition and normalization using precision, recall, and F1 at the mention (Men-) and document (Doc-) levels. Mention-level evaluation assesses individual annotated occurrences, whereas document-level evaluation assesses the normalized concepts identified within each article. Patient– phenotype association extraction was evaluated using micro- and macro-averaged precision, recall, and F1, together with the Jaccard similarity between predicted and reference phenotype sets. Micro-averaged metrics aggregate prediction counts, whereas macro-averaged metrics give equal weight to each evaluation unit. All results are reported as percentages.

## 4 Results

### 4.1 Corpus characteristics

PatientPheX comprises 200 full-text articles divided into training, development, and test sets in a 4:1:5 ratio, containing 80, 20, and 100 articles, respectively. The corpus includes 476 patients, 1,054 patient mentions, 55,980 biomedical entity mentions, and 7,359 patient-entity associations. Table 2 summarizes their distribution across the three subsets.

**Table 2.** Statistics of the PatientPheX dataset.

| Annotation | Train | Dev | Test | Overall |
| --- | --- | --- | --- | --- |
| Articles | 80 | 20 | 100 | 200 |
| Patients | 207 | 47 | 222 | 476 |
| Patient mentions | 435 | 99 | 520 | 1,054 |
| **Entities** | **22,495** | **5,465** | **28,020** | **55,980** |
| Phenotype | 6,040 | 1,565 | 7,580 | 15,185 |
| Disease | 7,724 | 1,978 | 9,064 | 18,766 |
| Gene | 4,151 | 1,033 | 5,374 | 10,558 |
| Variant | 1,617 | 316 | 1,985 | 3,918 |
| Chemical | 1,236 | 227 | 1,577 | 3,040 |
| Species | 1,662 | 344 | 2,285 | 4,291 |
| Cell line | 65 | 2 | 155 | 222 |
| Negated entities | 227 | 58 | 383 | 668 |
| Unnormalized entities | 493 | 138 | 774 | 1,405 |
| Composite entities | 1,660 | 303 | 2,099 | 4,062 |
| **Patient-entity relations** | **3,313** | **753** | **3,293** | **7,359** |
| Avg. patients / article | 2.59 | 2.35 | 2.22 | 2.38 |
| Avg. mentions / patient | 2.10 | 2.11 | 2.34 | 2.21 |
| Avg. phenotype relations/ patient | 8.32 | 8.51 | 7.75 | 8.07 |
| Avg. all relations / patient | 16.00 | 16.02 | 14.83 | 15.46 |

In the seven entity types, Disease was the most frequent entity type, with 18,766 mentions, followed by Phenotype (15,185) and Gene (10,558). Disease and Phenotype annotations overlapped substantially, reflecting expressions that can denote both a disease and a phenotypic abnormality, depending on context. The remaining annotations comprised 4,291 Species, 3,918 Variant, 3,040 Chemical, and 222 CellLine mentions. The corpus also included 668 negated mentions, 1,405 mentions without normalized identifiers, and 4,062 composite mentions, capturing additional complexity in clinical descriptions.

On average, each article contains an average of 2.38 patients, and each patient is referred to by 2.21 textual patient mentions. Each patient has an average of 15.46 biomedical entity associations, including 8.07 phenotype associations. The corresponding averages are broadly comparable across the training, development, and test sets, indicating similar patient and relation densities among the three subsets. These statistics highlight the document-level and patient-centered characteristics of the corpus, in which multiple patients and a large number of biomedical entities may need to be jointly resolved and associated within the same full-text article.

### 4.2 Entity recognition and normalization results

Table 3 presents the entity recognition and normalization results on the test set. The overall mention-level F1 was 87.31%, but performance varied substantially across entity types. Variant, Gene, and Disease achieved F1 scores above 90%, whereas Phenotype, Chemical, Species, and CellLine scored below 80%.

**Table 3.**
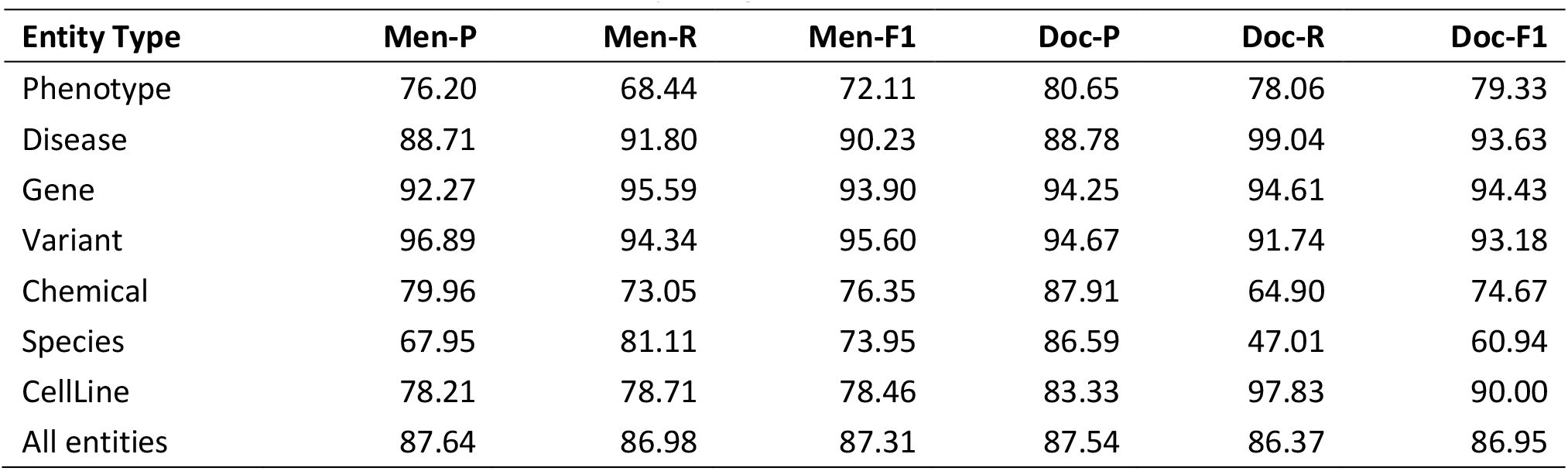
Performance of Entity recognition and normalization on the test set.

For Phenotype, recall was lower than precision, indicating incomplete recovery of annotated mentions. Document-level F1 increased to 79.33%, suggesting that recovering phenotype concepts within an article was more successful than identifying all their textual occurrences. This pattern is consistent with the variability of clinical descriptions, including composite expressions and wording that differs from canonical HPO labels.

Chemical and Species achieved mention-level F1 scores of 76.35% and 73.95%, respectively. Variation in chemical names and treatment descriptions may contribute to extraction errors. For Species, the inclusion of patient and family-member references extends the task beyond conventional organism names and may help explain the lower performance. Species document-level F1 was 60.94%, with recall of 47.01%, indicating substantial omissions under this evaluation. CellLine achieved mention-level and document-level F1 scores of 78.46% and 90.00%, respectively, although these results should be interpreted cautiously given its small test-set count of 155 mentions.

### 4.3 Patient-phenotype relation extraction results

We evaluated relation extraction under two conditions on the test set. The first used gold-standard entity annotations to assess patient attribution independently of upstream entity extraction errors. The second used phenotype candidates recognized and normalized by PhenoTagger to evaluate the combined extraction pipeline.

#### 4.3.1 Relation extraction with gold-standard entities

Table 4 shows the results for patient-phenotype RE with gold-standard entity annotations. GLM-5.1 achieved the strongest performance, with a micro-F1 of 88.04%, compared with approximately 71% for ATLOP and GLiM. Its advantage extended to macro-F1 and phenotype-set overlap, indicating that the improvement was not confined to the aggregate relation count.

**Table 4.** Performance of patient-phenotype relation extraction with gold-standard entity annotations.

| Model | Mic-P | Mic-R | Mic-F1 | Mac-P | Mac-R | Mac-F1 | Jaccard |
| --- | --- | --- | --- | --- | --- | --- | --- |
| Qwen3-8B | 74.06 | 84.27 | 78.84 | 67.86 | 77.98 | 69.23 | 61.20 |
| GLM-5.1 | 84.89 | 91.43 | 88.04 | 84.13 | 89.65 | 84.14 | 78.08 |
| GPT-5.4-mini | 74.90 | 90.08 | 81.79 | 75.69 | 85.96 | 76.85 | 69.14 |
| ATLOP | 64.92 | 79.10 | 71.31 | 60.91 | 75.79 | 63.78 | 53.00 |
| GLiM | 66.50 | 77.95 | 71.77 | 66.96 | 74.29 | 65.27 | 55.06 |

All methods achieved higher recall than precision, indicating that false-positive associations remained an important source of error even with correct entity annotations. GLM-5.1 combined high recall with the highest precision, whereas GPT-5.4-mini achieved similar recall but lower precision. This distinction suggests that GLM-5.1 was more effective at excluding unsupported associations while retaining relevant findings. ATLOP and GLiM had similar micro-F1 scores, although GLiM achieved higher macro-F1 and Jaccard similarity. These results show that correct entity annotations alone do not ensure accurate patient attribution.

Resolving patient references and distinguishing patient-specific findings from information about other individuals or general background remain important challenges. Nevertheless, attributing the observed differences specifically to coreference resolution, negation handling, or long-range reasoning would require targeted error analysis.

#### 4.3.2 Pipeline relation extraction with predicted entities

Table 5 reports the final patient-phenotype association results when PhenoTagger predictions were supplied to the relation extraction methods. GLM-5.1 remained the strongest model across all reported metrics, but its micro-F1 decreased from 88.04% to 62.97%. The range of micro-F1 scores across methods also narrowed substantially, from 71.31–88.04% with gold-standard entities to 56.50–62.97% with predicted entities. Thus, the advantages observed in patient attribution with correct inputs were substantially reduced in the complete extraction pipeline.

**Table 5.** Pipeline performance of patient–phenotype relation extraction using PhenoTagger predictions.

| Model | Mic-P | Mic-R | Mic-F1 | Mac-P | Mac-R | Mac-F1 | Jaccard |
| --- | --- | --- | --- | --- | --- | --- | --- |
| Qwen3-8B | 55.23 | 62.73 | 58.74 | 51.60 | 58.36 | 51.88 | 39.77 |
| GLM-5.1 | 63.98 | 62.00 | 62.97 | 57.75 | 57.08 | 54.98 | 43.93 |
| GPT-5.4-mini | 59.98 | 61.26 | 60.62 | 54.84 | 56.29 | 53.14 | 42.03 |
| ATLOP | 52.76 | 60.89 | 56.53 | 48.44 | 57.17 | 49.14 | 36.74 |
| GLiM | 53.89 | 59.38 | 56.50 | 53.60 | 54.45 | 49.43 | 37.95 |

Micro-recall converged to a relatively narrow range across models, while precision varied more widely. GLM-5.1’s leading F1 therefore reflected its higher precision rather than the highest recall. Qwen3-8B recovered slightly more reference associations but also produced more false positives. ATLOP and GLiM performed similarly and remained below the LLM-based methods.

The decline across all methods is consistent with the propagation of upstream recognition and normalization errors. When relation prediction is restricted to supplied candidates, phenotype concepts missing from the candidate set cannot be recovered by the relation extraction stage. Incorrectly recognized or normalized candidates can also lead to invalid associations. These effects help explain why strong performance with gold-standard entities did not translate directly into comparable pipeline performance.

Jaccard similarities ranged from 36.74% to 43.93%, showing limited overlap between the predicted and reference phenotype sets. The pipeline therefore remains insufficient for reliably reconstructing complete patient-specific profiles, even when it identifies many individual associations correctly.

### 4.3 Discussion

The benchmark results reveal two major challenges of information extraction from full-text patient-centered biomedical literature. The first is the linguistic complexity of clinical phenotype descriptions. Although several conventional biomedical entity types can already be recognized with relatively high accuracy, phenotype extraction remains considerably more difficult. Phenotypes are often expressed through free-form clinical descriptions rather than standardized terminology, and a single clinical expression may contain several related abnormalities or require contextual interpretation before it can be mapped to an HPO concept. Consequently, errors may arise from both mention boundary detection and concept normalization. The difference between mention-level and document-level performance further suggests that identifying the correct phenotype concept is sometimes easier than locating its exact textual realization.

The second challenge is patient attribution. Articles may describe several individuals using names, pronouns, family relationships, or pedigree identifiers, while their findings are distributed across sections. Accurate extraction requires linking these references and distinguishing findings for the target patient from those for relatives, controls, or previously reported cases. Long-range context further increases the difficulty of patient-phenotype association extraction. Patient mentions and their associated phenotypic findings are not always located in the same sentence or paragraph. A patient may first be introduced in the case presentation, while relevant phenotypic findings are described later in physical examination, laboratory, imaging, or follow-up sections. This suggests that effectively integrating document-level context and maintaining patient-specific information across long documents are important for this task.

The pipeline results further show that entity extraction and patient attribution must be evaluated together. GLM-5.1 performed best in both conditions, but its substantial decline with predicted entities demonstrates that improved attribution alone does not eliminate upstream limitations. This finding motivates methods that can reconsider uncertain phenotype candidates using patient context, while preserving reliable links between textual evidence, normalized concepts, and patient identities.

Negation and composite expressions also require careful representation. An absent phenotype should remain distinguishable from an observed finding, and a composite expression may need to be resolved into several concepts before patient attribution. PatientPheX provides annotations for investigating these challenges, but the current aggregate results do not quantify performance on these specific subsets. Future analyses stratified by negation, composite mentions, number of patients, and distance between supporting mentions would clarify which features most strongly affect extraction accuracy.

Overall, PatientPheX supports complementary evaluation of entity extraction, patient attribution with gold-standard inputs, and the combined pipeline. The differences between these settings highlight the need to improve both candidate quality and contextual attribution to produce more complete and accurate patient-specific phenotype profiles.

## 5 Conclusion

In this work, we presented PatientPheX, a corpus of 200 full-text biomedical articles for patient-specific phenotype information extraction. Corpus construction combined automated pre-annotation, LLM-assisted verification, independent manual annotation, and adjudication. The corpus integrates annotations of seven biomedical entity types, entity normalization, negated findings, and composite phenotype mentions with explicit patient-entity associations across sentences, paragraphs, and sections. These annotations support the development and evaluation of methods that organize dispersed biomedical findings into structured patient-specific profiles. Benchmark experiments evaluated entity recognition and normalization alongside patient attribution under both gold-standard and predicted entity conditions, highlighting the importance of assessing the complete extraction pipeline.

The current corpus is limited in size and primarily comprises case reports and case series, which may restrict generalization to other biomedical text types. Pipeline performance remains constrained by errors in phenotype recognition, normalization, and patient attribution. The results indicate room for improvement in identifying phenotype expressions and assigning findings to the correct patients across full-text documents. Future work will expand the corpus and its coverage, and investigate methods that jointly model phenotype semantics, patient identity, and document-level context. Further studies will assess the utility of extracted patient profiles in phenotype-driven biomedical research.

## Acknowledgements

This work was supported by the National Natural Science Foundation of China (No. 62302076, 62276043, 62676065) and the Fundamental Research Funds for the Central Universities (No. DUT25YG108). The authors would like to thank Jiewei Qi, Pengqi Zheng, and Xiwen Bi for their contribution with data curation.

## Notes

### Competing Interest Statement

The authors have declared no competing interest.

